# A proof-of-principle study of the effect of combined haloperidol and levodopa administration on working memory-related brain activation in humans

**DOI:** 10.1101/219436

**Authors:** Peter Van Ruitenbeek, Dennis Hernaus, Mitul Ashok Mehta

## Abstract

**Background and Purpose:** Cognitive deficits including impaired working memory are a hallmark feature of schizophrenia. Changes in prefrontal cortex function modulated by dopamine D1 receptors, play a potentially important role in the pathology underlying such deficits. However, pharmacological interventions that selectively engage the D1 receptor are severely restricted for research in humans. The present study is a proof-of-principle for enhancing cognitive performance and associated brain activation via indirect D1 stimulation. Here, we combine the non-selective dopamine agonist L-dopa with the D2-antagonist haloperidol, theoretically producing increased stimulation at the D1 receptor.

**Experimental Approach:** Fourteen healthy volunteers received placebo or combined carbidopa (125 mg, 100mg L-dopa) plus haloperidol (2 mg) orally on two separate occasions according to a within-subjects cross-over design. Drug-induced differences in brain activity were assessed during an N-back working memory task in a 3T magnetic resonance imaging environment.

**Key Results:** Drug treatment was associated with a reduction in activity in a large number of brain areas, most prominently occipital/temporal brain areas during 2-back performance, which may be due to the effects of haloperidol specifically. Drug treatment was also associated with greater functional connectivity within parts of the salience network during all N-back trials.

**Conclusion and Implications:** This preliminary study provides initial evidence for combined L-dopa/haloperidol modulation in cognition-related brain areas and networks, which is relevant for the treatment of cognitive impairments in mental illness.

## INTRODUCTION

Schizophrenia is characterised by broad and persistent cognitive deficits, including impaired working memory and episodic memory, executive functioning and attention (Fioravanti *et al*., 2012). While currently available pharmacological treatments are primarily aimed at decreasing positive symptoms (e.g. hallucinations), they do not alleviate cognitive deficits (Marder, 2006). Cognition-enhancing agents for schizophrenia represent a core unmet need: in addition to direct treatment of cognitive deficits, these agents may promote functional independence via improved insights into disease and therapy (Green *et al*., 2000). Here, we aim to provide proof-of-principle evidence for a potential cognition-enhancing treatment, which could inform a novel treatment strategy for individuals with schizophrenia (Saha *et al*., 2007).

Dopamine D2 receptor antagonists are the most widely-used class of pharmacological agents in schizophrenia. It is thought that D2 receptor blockade, primarily in striatum, is the main mechanism-of-action by which antipsychotics decrease positive symptoms. Importantly, however, D2 receptor blockade does not appear to explain the modest improvement in cognitive impairments seen in some patients (Goldberg *et al*., 2007) and treatment with second-generation antipsychotics does not improve working memory and attentional functions (Nielsen *et al*., 2015), underlining the need for alternative approaches.

Preclinical research has established that dopamine D1 receptors essentially modulate prefrontal cortex (PFC)-mediated working memory (Sawaguchi *et al*., 1991). For example, D1 agonism improves working memory in aged monkeys (Castner *et al*., 2004), while D1 antagonism negatively impacts spatial working memory abilities. At the neural level, D1 receptor activity increases signal-to-noise-ratio (SNR) in neural networks, including PFC, most likely by decreasing spontaneous firing of neurons (Seamans *et al*., 2001). An optimal level of PFC dopamine activity has been described by many studies in experimental animals and humans, with insufficient and excessive D1 receptor activation leading to reduced SNR (Akaike *et al*., 1987), and an inverted-U shaped relationship between dopamine activity and cognitive performance (Cools *et al*., 2011).

Cognitive deficits in schizophrenia may be the consequence of altered dopamine D1, rather than D2, function in PFC (Abi-Dargham *et al*., 2002). This notion is supported by the observation that D1 receptor density in PFC correlates with cognitive performance in schizophrenia (Abi-Dargham, 2003) and schizotypal personality disorder (Thompson *et al*., 2014). Moreover, in animal models of schizophrenia, D1 receptor agonism can reverse cognitive impairments (McLean *et al*., 2009). In light of this evidence, D1 receptors have long been considered potential treatment targets (Goldman-Rakic *et al*., 2004).

Dopamine D1 receptors can be stimulated directly or indirectly, although currently available compounds have significant drawbacks (Arnsten *et al*., 2016). For example, amphetamine non-selectively enhances catecholaminergic and serotonergic activity, in addition to abuse potential. The D1receptor agonist SFK-38393 is potentially useful, as it reverses phencyclidine-induced cognitive deficits in experimental animals, but it has not been used in humans (McLean *et al*., 2009). DAR-0100A is the only selective D1 receptor agonist available for use in humans, although it has exclusively been used at doses that do not produce measurable D1 receptor occupancy (Slifstein *et al*., 2011) and produces a range of side effects (George *et al*., 2007).

While there were preliminary findings of improved spatial working memory in schizotypal personality disorder (Rosell *et al*., 2015), DAR-0100A did not improve executive functions or cognition-related brain function in schizophrenia (Girgis *et al*., 2016). This may be related to the agent’s short half-life (Blanchet *et al*., 1998), complicating successful treatment. An alternative and accessible approach is to increase dopamine turnover nonselectively and simultaneously block D2 receptors, hypothetically producing dopamine D1 receptor agonism. Dopamine turnover can be increased non-selectively by administration of its precursor L-dopa, which increases dopamine synthesis in the central nervous system and periphery (Rosen *et al*., 1986). Peripheral increases in dopamine can be blocked by carbidopa, further increasing central dopamine availability (Rosen *et al*., 1986). Haloperidol has a mixed profile, but acts principally as a dopamine D2-antagonist.

In the present proof-of-principle study, we hypothesised that simultaneous administration of L-dopa/carbidopa and haloperidol would decrease PFC brain activity and increase functional connectivity (Meyer-Lindenberg *et al*., 2001) during a working memory paradigm. Here, drug-induced *decreases* in brain activity in combination with functional connectivity *increases* indicate more efficient network activity, the result of increased SNR (Callicott *et al*., 2000). As drug-induced performance increases may be difficult to establish in healthy volunteers, who already function at a near-optimal level, we relied on the sensitive nature of BOLD fMRI during N-back task performance. The N-back is a well-established paradigm for probing working memory function, has been used extensively in schizophrenia [e.g. (Bertolino *et al*., 2003)] and is sensitive to dopaminergic drug effects (Mattay *et al*., 2000).

## Methods

### Participants

We recruited fourteen healthy right-handed male volunteers aged between 19 and 38 years (mean±sd = 25.0±5.1 years) by circular e-mail sent to students and staff from King’s College London and were financially compensated for their time. Using BOLD for the DLPFC from an in-house N-back dataset, we estimated that with 14 participants we would have a power of ∼0.55 to detecting a 50% reduction in BOLD signal *amplitude* in the drug condition at p<0.05. Inferences were planned for cluster correction statistics for which *a-priori* power analyses is difficult. Participants’ physical and mental status were assessed by a screening involving a urine analysis and test for the presence of drugs of abuse (amphetamines, methamphetamines, THC, methadone, opiates, phencyclidine, barbiturates, benzodiazepine, tricyclic antidepressants), a 12-lead electrocardiogram, measurements of heart rate and blood pressure, breath alcohol concentration and blood chemistry and haematology.

Volunteers were excluded if they showed any evidence or history of clinically significant renal, pulmonary, gastrointestinal, cardiovascular, hepatic, psychiatric or neurological disease/disorder, including epilepsy or seizures and more than one febrile convulsion. In addition, volunteers were excluded from the study if they used any prescribed or non-prescribed drugs except paracetamol and acetaminophen, drank more than 28 standard alcohol units per week, smoked more than 5 cigarettes per day, were treated with a new chemical entity within the past 3 months, had a known sensitivity to any of the study medications, a Body Mass Index outside the limits of 18-30 kg.m^−2^, non-removable metallic items in/on their body or signs of claustrophobia.

All participants gave written informed consent before they entered the study that was approved by the King’s College Research Ethics Committee (RECnr.: 10/H0807/13) and was conducted in accordance with the World Medical Association Declaration of Helsinki and its amendments (World-Medical-Association, 1964, 1996, 2008, 2013).

### Experimental design and treatment

The study was a two-way double blind, placebo-controlled design. Study medications were combined oral doses of haloperidol 2mg and L-dopa 100mg/carbidopa 25mg or placebo (ascorbic acid) administered according to a double dummy procedure and administration order was randomised and counterbalanced across participants. Haloperidol 2 mg has been shown to block 60-75% of D2 receptors and doses up to 3mg are generally well-tolerated (Legangneux *et al*., 2000). L-dopa 100mg/carbidopa 25mg is also well-tolerated and produces minimal side effects in healthy volunteers (Floel *et al*., 2008). However, nausea occasionally occurs, which we aimed to prevent by administration of peripherally-acting dopamine antagonist domperidone, which was administered prior to the start of the experiment on both study days (drug *and* placebo).

### Procedure

Volunteers visited the Centre for Neuroimaging Sciences three times. The first visit was a screening/training visit during which volunteers gave their written informed consent, were medically screened and performed the tasks in a mock scanner to familiarise them with the MRI environment and task procedures. The second and third visits were scanning visits. After confirming suitability (including a physical assessment conducted by a physician), subjective mood was assessed using 16 visual analogue scales (Bond *et al*., 1974). Volunteers received haloperidol 2mg and domperidone or placebo at time T0. After waiting for 60 minutes, they received L-dopa 100mg/carbidopa 25mg or placebo (T60), ensuring that plasma peak levels of both drugs occurred at similar times [T_max_ Haloperidol = 1.7-6.h (Kudo *et al*., 1999), T_max_ L-dopa = 15-60 min (Contin *et al*., 2010)]. 120 Minutes after the administration of the first dose (T120) a second assessment of subjective mood was performed using a visual analogue scale. Volunteers received a small standardised lunch and entered the scanner 150 minutes after the first dose (T150). Visual stimuli were back-projected on a screen, which the volunteer could see using periscopic mirrors. After scanning, volunteers’ subjective mood was assessed again. Finally, a physical examination was performed as part of a discharge assessment by the physician.

### Materials and tests

#### N-back task

The N-Back task is a well-established working memory task that reliably activates the dorsolateral PFC (Owen *et al*., 2005) and has been used in many studies to assess drug-induced changes in working memory performance The task consisted of four conditions: 1-back, 2-back, 3-back and a 0-back control condition. Each task condition occurred three times as blocks of 14 letters sequentially presented on a screen for 2 seconds per letter. Before each 28-second block of trials, the upcoming condition was briefly presented for 2 seconds. Volunteers responded to cues with a button press on a response box using their right index (target) and left index (non-target) finger. A target stimulus was defined as a letter that matched the previous one in the 1-back condition (e.g. ‘A’ followed by ‘<u>A</u>’), a letter presented two letters earlier in the 2-back condition (e.g. ‘A’, ‘B’, ‘<u>A</u>’), a letter presented three letters earlier in the 3-back condition (e.g. ‘A’, ‘B’, ‘C’, ‘<u>A</u>’) or was an X in the 0-back condition (e.g. ‘A’, ‘B’, ‘<u>X</u>’). The total length of the task was 6 minutes and 20 seconds. Reaction time and responses were recorded and average reaction time and number of correct responses were dependent variables.

#### Subjective mood ratings

Subjective evaluations of alertness, contentedness, and calmness were assessed using a series of 16 visual analogue scales (100 mm), which provided factor analytically defined summary scores for ‘alertness’, ‘contentedness’, and ‘calmness’ (Bond *et al*., 1974). Participants were asked to indicate their current mood state by marking a horizontal line in between two extremes of a given mood dimension, e.g. alert-drowsy.

### Image acquisition

All MRI data were collected at the Centre for Neuroimaging Sciences, King’s College London using a General Electric 3 Tesla Signa HDx scanner (General Electric, Milwaukee) with an 8 channel head coil. Volunteers lay in a supine head first position in the scanner. One hundred and eighty-six T2*-weighted images were acquired during N-back task performance using a gradient echo planar imaging sequence with repetition time/echo time=2000/30 msec, flip angle=75°, 37 slices (sequential, top-bottom), slice thickness/gap=3/0.3 mm, in-plane resolution 3.3 mm2, and field of view 21.1 cm. To allow for accurate normalisation to a standard space a whole-brain three-dimensional inversion recovery prepared spoiled gradient echo (IR-SPGR) scan, was also collected with isotropic 1.1-mm voxels in a scan time of approximately 6 min (repetition time/echo time 6.96/2.82 msec; inversion time = 450 msec; excitation flip angle = 20°).

### Image pre-processing

Imaging data were analysed in SPM8 (http://www.fil.ion.ucl.ac.uk/spm/) using Matlab (V7.0.1.) (https://mathworks.com) on a UNIX platform. Slices were corrected for acquisition time using the middle slice as reference. The data were corrected for translational and rotational movement in 3 dimensions, first by registering all images to the first in the series and then to the mean image. In all runs, volunteer head movement did not exceed the limit of 1 voxel on no more than 3 occasions within one run, either translational or rotational (i.e. a rotation equating to 1 voxel at the brain surface). The functional data were co-registered with the high resolution T1-weighted image data using the mean image to determine the parameters. The T1-weighted images were normalised to standard MNI space using unified segmentation with the parameters applied to the functional time series which were then smoothed using a Gaussian kernel of 8mm of full width at half maximum. All scans were visually inspected for quality of pre-processing.

### Data and statistical analysis

#### Behavioural data

The data and statistical analysis comply with the recommendations on experimental design and analysis in pharmacology (Curtis *et al*., 2015). Two participants did not perform above chance level on 2-back and 3-back trials and their data were excluded from analysis. For the remaining 12 participants a 2x4 General Linear Model (GLM) for Repeated Measures analyses were performed for reaction time and correct responses, with Treatment (2 levels: haloperidol/L-dopa, placebo) and Level of difficulty (4 levels: 0-, 1-, 2- and 3-back) as within subject factors. We tested for a main effect of Treatment and Level of difficulty and their interaction at α=.05 significance level, corrected using Greenhouse-Geisser method when the sphericity assumption was violated. A significant main effect of Level of difficulty was further specified using contrasts between 0-back condition and 1-, 2-, 3-back condition. A significant interaction between Level of difficulty and Treatment was further specified using drug-placebo contrasts within each difficulty level.

Visual analogue scales data from all 14 participants were analysed separately for three factors (Alertness, Contentedness and Calmness) using GLM for repeated measures with Treatment (2 levels: haloperidol/L-dopa, placebo), and Time Point (3 levels: T0, T120, and T210) as within subjects factors. Significance level was set at α=.05, and was corrected using Greenhouse-Geisser when the sphericity assumption was violated. All behavioural data were analysed using SPSS 18 (SPSS inc., 2009).

#### Imaging data

For 12 participants performing above chance level, functional images were first analysed at the single-subject level in the framework of the General Linear Model. The design matrix comprised 11 regressors; 0-, 1-, 2-3-back trial blocks, visually presented instructions, and 3 translational and 3 rotational movement parameters, for which Beta weights were estimated. Next, contrasts were defined between 0-back and 1-, 2-, 3-back, representing activation during 0-back subtracted from activation during the other N-back difficulty levels. For the 2^nd^ level analysis all first level contrasts were resliced to match the FMRIB Software Library’s [FSL; (Jenkinson *et al*., 2012)] MNI152 2mm template. Next, all contrasts images were concatenated into a single 4D file. Using FSL’s randomise (Winkler *et al*., 2014), F-contrasts for Treatment (2 levels: haloperidol/L-dopa, placebo) and Level of difficulty (3 levels: 1-, 2-, 3-back > 0-back contrasts) and their interaction were calculated in a model with Treatment and Level of difficulty as fixed factors and subject as random factor. Significant (TFCE, p<.05 FWE corrected) main effects or interaction were further analysed using t-contrasts.

In addition, a generalized Psycho-Physiological Interaction (gPPI: (McLaren *et al*., 2012) analysis was performed to determine if combined haloperidol/L-dopa treatment affected task-dependent correlations between brain areas within the network activated during N-back task performance. First, the dorsolateral prefrontal cortex (DLPFC) was chosen as an *a-priori* seed region given its prominent role in N-back performance (Owen *et al*., 2005) and its involvement in schizophrenia working memory pathology (Abi-Dargham *et al*., 2002). The region was identified in the current data set by performing an F-test for the main effect of Level of difficulty taken from the analysis described above. Peak MNI coordinates from this analysis in the current data set [i.e. right DLPFC: 44, 26, 36] were used to guide the identification of active brain areas in all runs separately, locating the nearest local maximum. Subsequently, a six mm sphere was created around the peak voxel that was masked with the activation map from the F-test for Level of difficulty resulting in a volume of interest (VOI) containing only active voxels per run. The VOIs were fed into the gPPi analysis using the SPM toolbox provided by McLaren *et al*. (2012) (http://www.nitrc.org/projects/gppi). gPPI has the advantage over PPI that it accommodates multiple task conditions, that may explain variance of the dependent variable, and provides a better model fit (McLaren *et al*., 2012) Using the gPPi toolbox, contrast images were created representing memory load dependent activation correlated with activation of the seed region (PPI term). Finally, a 2^nd^ level analysis was performed to determine Treatment differences in PPI terms: i.e. differential effects of haloperidol/L-dopa and placebo on memory load-dependent DLPFC functional connectivity. Using permutation testing with FSL’s randomise, the F-contrasts for main effect of Treatment was calculated within the N-back network. To explore the direction of findings from the F-test, t-contrasts were conducted using TFCE and a p<.05, FWE corrected.

## RESULTS

### Behavioural data

#### N-back task

Increased level of difficulty was accompanied by a significantly lower percentage of correct responses and significantly increased the average reaction time. Compared with the 0-back condition, responses were significantly slower during 1-back blocks, and significantly fewer correct- and slower responses were given during 2-back blocks and 3-back blocks.

The combination of haloperidol and L-dopa did not significantly affect the percentage of correct responses or the average reaction time on any level of difficulty or across levels of difficulty. However, a trend for more correct responses after haloperidol and L-dopa was observed for 0-back (t_11_=-1.865, p<.089) and 1-back task (t_11_=-1.820, p<.096) task conditions, in absence of a significant increase in reaction time. Please see table 1 for descriptive and significance statistics.

**Table 1.**
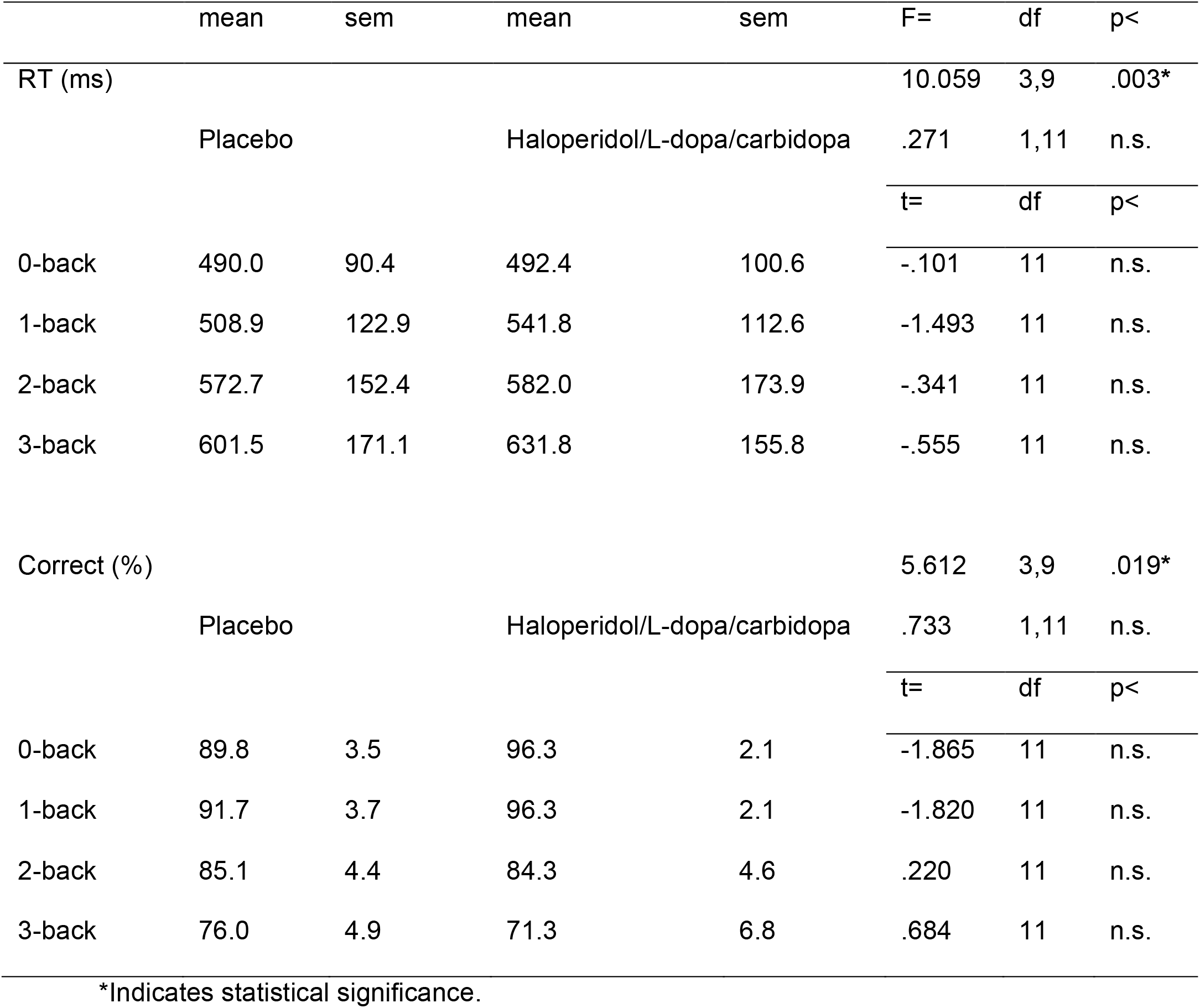
Descriptive statistics and test of significance of reaction times and % correct for the 4 levels of difficulty of the N-back task, performed after a single dose of placebo or haloperidol/L-dopa/carbidopa drug treatment. Performance on both measures was significantly affected by level of difficulty. Drug treatment did not significantly affect either measure. RT = reaction time, sem = standard error of measurement, df = degrees of freedom, n.s. = not significant.

#### Subjective Mood ratings

Time Point (TP) showed a significant effect on measures of alertness and contentedness. Alertness significantly decreased over time points and contentedness was lower at T120 compared with T0 and with T210. Drug treatment did not affect subjective ratings of alertness or contentedness. The treatment by TP interaction showed a trend, driven by haloperidol/L-dopa increasing calmness over time compared with placebo. Contrasts between haloperidol/L-dopa and placebo were significant or close to significance for differences between calmness measured at T210 and T0, and T210 and T120.

### Imaging data

#### N-back: Level of difficulty effects

F-contrast for the main effect of Level of difficulty indicating any activation differences with 0-back resulted in a large number of significant clusters of activation (see figure 2). In addition, t-contrasts assessing where brain activity increased with task difficulty revealed a number of significant areas in the 2-back > 1-back contrast which closely resembled the activation pattern of the general F-contrast (figure 2). In addition, the 3-back>2-back t-contrast did not reveal any significant activity differences (see figure 2).

**Figure 1.**
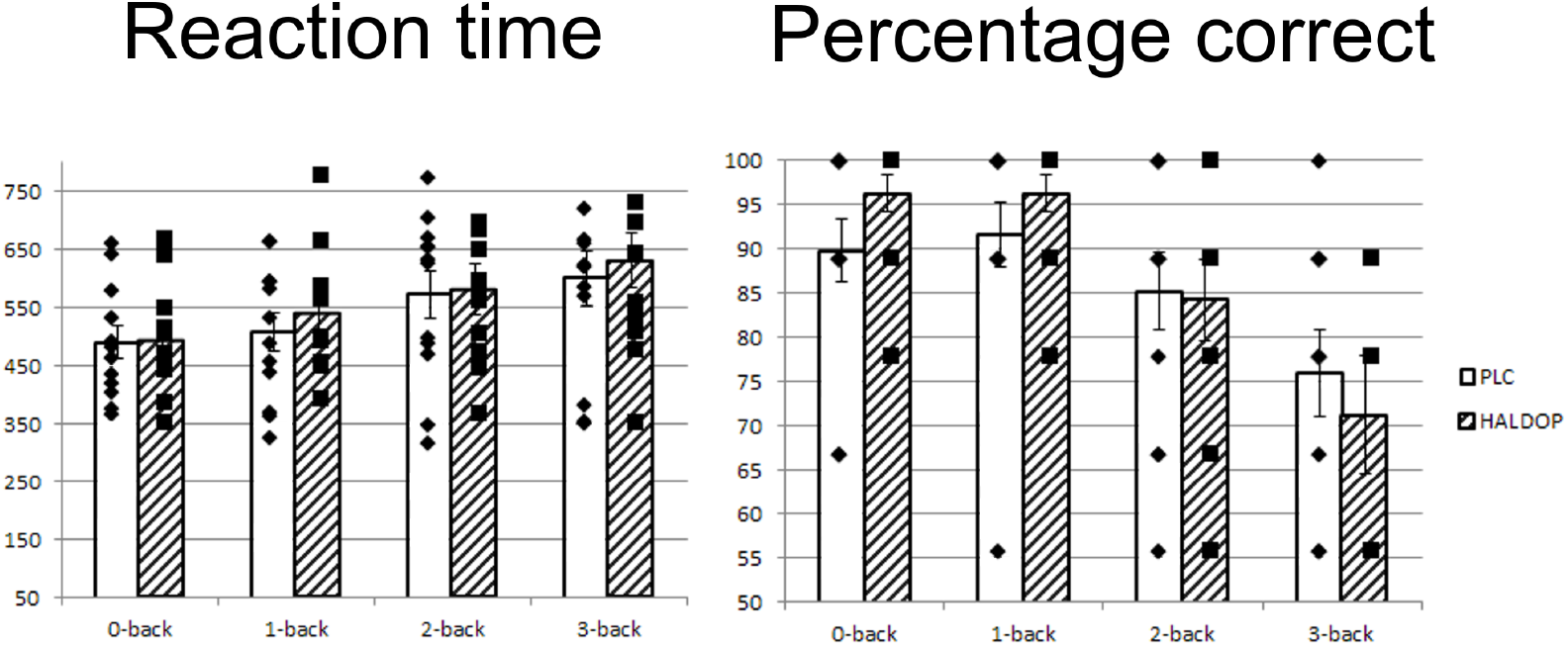
Performance on different levels of N-back difficulty after receiving placebo or haloperidol+L-dopa (n=12). No significant drug effects were observed, although drug treatment was followed by more accurate responses during 0-back and 1-back at a trend level (p<.1).

**Figure 2.**
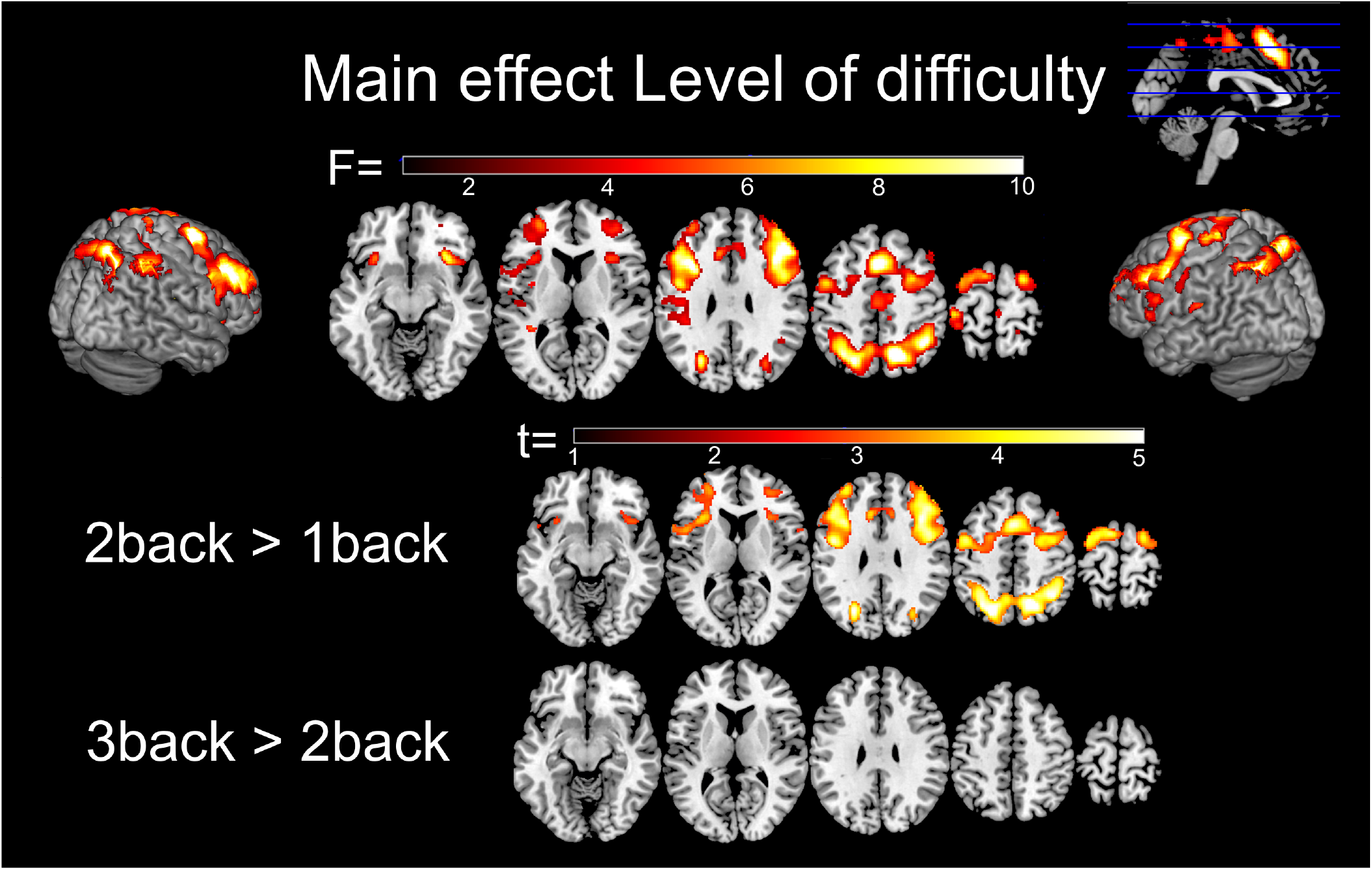
Main effects of Level of difficulty (within subject variable, n=12) on brain activation, as well as 2-back>1-back and 3-back>2-back contrasts. The 2-back condition evoked activity most robustly in premotor cortex (BA6), supplementary motor area (BA6), anterior cingulate (BA32), superior parietal lobule (BA7), inferior parietal lobule (BA40), Broca’s area (BA44, 45), middle frontal gyrus (BA46), insula (BA47, 48), frontal pole (BA10).

#### N-back: Treatment effect

An F-test for Treatment effects indicated significant activation differences between drug and placebo sessions. T-contrasts revealed that combined haloperidol/L-dopa administration significantly decreased brain activation in a wide range of regions compared to placebo (see figure 3 and table 2).

**Figure 3.**
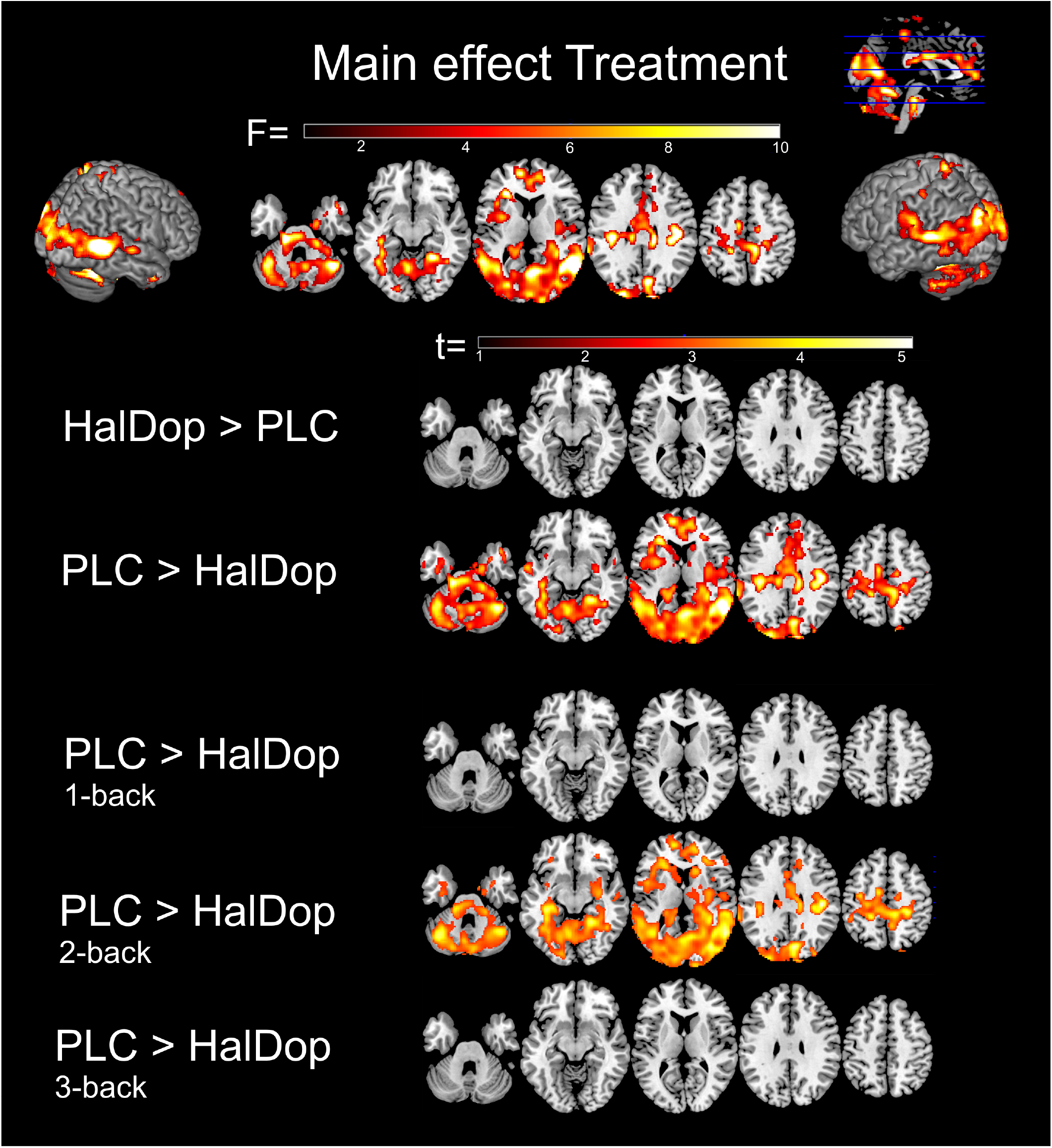
A main effect of Treatment (within subject variable, n=12) on brain activation during N-back performance was observed in a large number of brain areas. Specific t-contrasts indicate that brain activation decreased after drug treatment, specifically during the 2-back task condition.

**Table 2.** Brain areas showing less activation following haloperidol/L-dopa compared with placebo across all N-back conditions. Brodmann area (BA), peak t-values, peak p-values and MNI coordinates indicate peak cluster voxels.

| General Area | Specific Area | BA | Peak p-value<br>(FWE) | Peak t-value | Peak coordinates |  |  |
| --- | --- | --- | --- | --- | --- | --- | --- |
|  |  |  |  |  | X | Y | Z |
| Frontal cortex | R. Frontal Pole | 10 | 0.0429 | 3.03 | 16 | 70 | 4 |
|  | R. Frontal Pole | 10 | 0.043 | 3.01 | 10 | 68 | 2 |
| Sensorimotor areas | R. Postcentral Gyrus | 4 | 0.0019 | 5.02 | 6 | -36 | 54 |
|  | L. Primary | 3b | 0.003 | 4.87 | -18 | -42 | 52 |
|  | R. Premotor Cortex | 6 | 0.0284 | 4.62 | 56 | -2 | 44 |
|  | R. Primary | 1 | 0.0345 | 3.53 | 56 | -12 | 44 |
|  | R. Secondary | 48 | 0.0009 | 6.7 | 42 | -20 | 28 |
| Occipital cortex | L. Occipital Pole | 18 | 0.0011 | 4.82 | -18 | -94 | 34 |
|  | R. Occipital pole | 18 | 0.0012 | 5.24 | 10 | -94 | 32 |
|  | R. Visual Cortex V1 | 17 | 0.0006 | 4.87 | 6 | -66 | 10 |
|  | R. Visual Cortex V2 | 18 | 0.0006 | 4.92 | 8 | -78 | 28 |
|  |  | 18 | 0.0006 | 4.91 | 10 | -72 | 20 |
|  |  | 18 | 0.0006 | 4.9 | 10 | -62 | 2 |
|  | R. Visual Cortex V5 | 37 | 0.0006 | 4.92 | 40 | -62 | 10 |
|  | L. Occipital Pole | 18 | 0.001 | 5.14 | -26 | -94 | 26 |
| Temporal/Occipital cortex | R. Temporal Occipital | 14 | 0.0006 | 4.97 | 26 | -48 | -14 |
|  | R. Temporal Occipital | 37 | 0.0006 | 4.97 | 22 | -46 | -16 |
|  | R. Temporal Occipital | 18 | 0.001 | 4.9 | 42 | -46 | -18 |
|  | L. Angular Gyrus | 37 | 0.0008 | 4.93 | -58 | -56 | 12 |
|  | R. Angular gyrus | 37 | 0.0002 | 7.11 | 56 | -60 | 8 |
| Cerebellum | L. Left Cerebellum V |  | 0.0008 | 4.85 | -4 | -58 | -14 |
|  | L. Cerebellum VI |  | 0.0009 | 4.82 | -36 | -56 | -26 |
|  | R. Cerebellum VI |  | 0.0006 | 5.06 | 12 | -66 | -22 |
|  | R. Cerebellum Crus I |  | 0.0006 | 5.04 | 42 | -60 | -28 |
| Subcortical | L. Thalamus |  | 0.0275 | 2.7 | -2 | -10 | 2 |
|  | Brain Stem |  | 0.001 | 5 | -6 | -24 | -26 |
|  | Brain stem |  | 0.0009 | 5.54 | -18 | -26 | -32 |

Although no significant Treatment by Level of difficulty interaction was observed, planned t-contrasts did reveal a significant Treatment effect during the 2-back condition. During the 2-back condition, combined haloperidol/L-dopa administration reduced activity in a large number of clusters that mostly overlapped with the clusters from the overall F-contrast for Treatment. A noticeable exception was reduced activation in bilateral caudate, right amygdala, and right thalamus; see figure 3 and table 3.

**Table 3.**
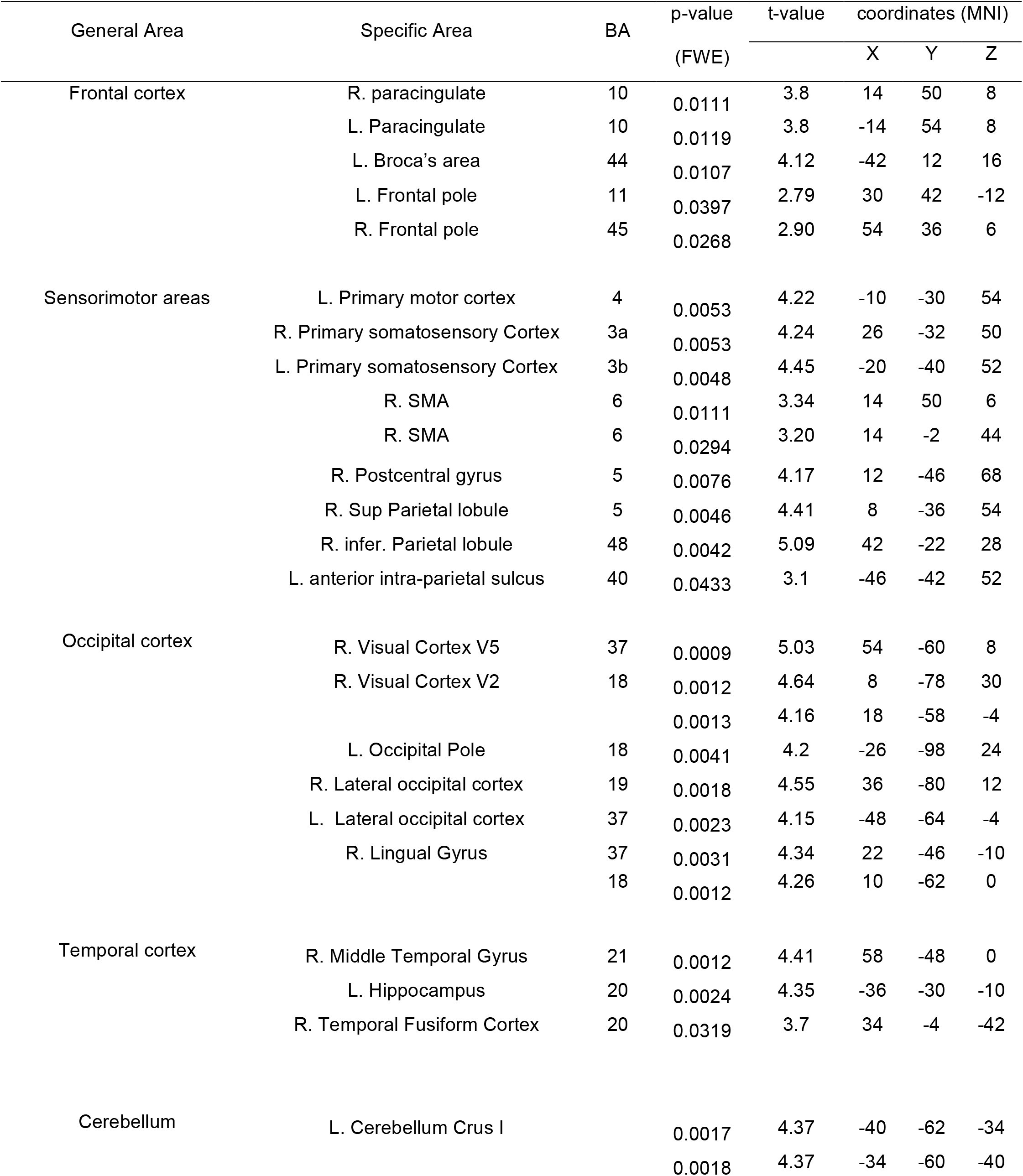
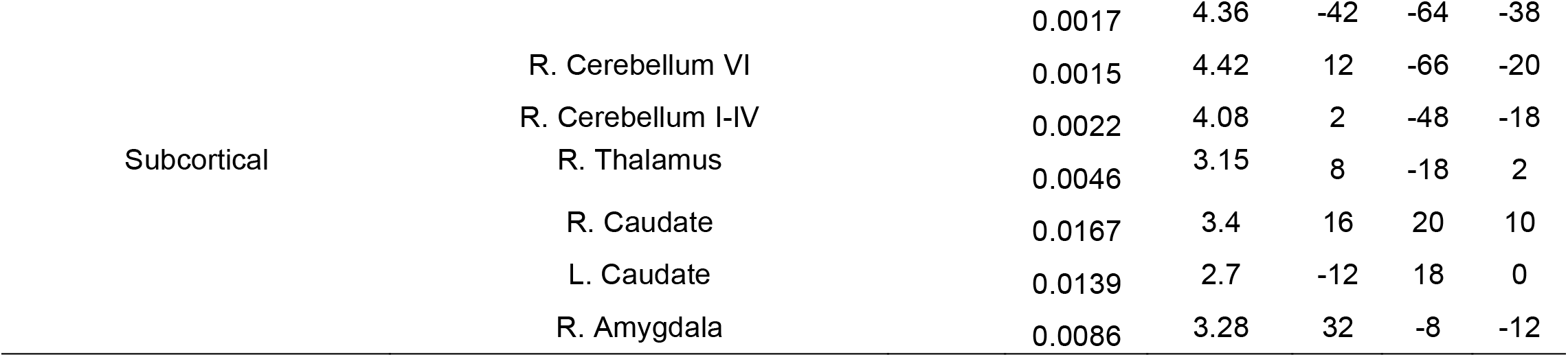
Brain areas in which haloperidol/L-dopa produced less activation compared with placebo during the 2-back condition. Brodmann area (BA) and peak t-values, peak p-values and MNI coordinates indicate peak voxels within the clusters.

#### N-back connectivity

Planned exploratory t-tests showed that combined haloperidol/L-dopa treatment increased task-dependent connectivity between DLPFC and several clusters for all n-back contrasts (1-, 2-, and 3-back vs. 0-back). The first cluster included right superior frontal gyrus (peak voxel at BA8), bilateral paracingulate gyrus (peak voxel at BA32), and right middle frontal gyrus (peak voxel at BA45). The second cluster included two peaks in the right premotor cortex (peak voxels at BA6). Finally, one cluster was detected in the left premotor cortex (peak voxel at BA6). Please see figure 4 and table 4 for details.

**Figure 4.**
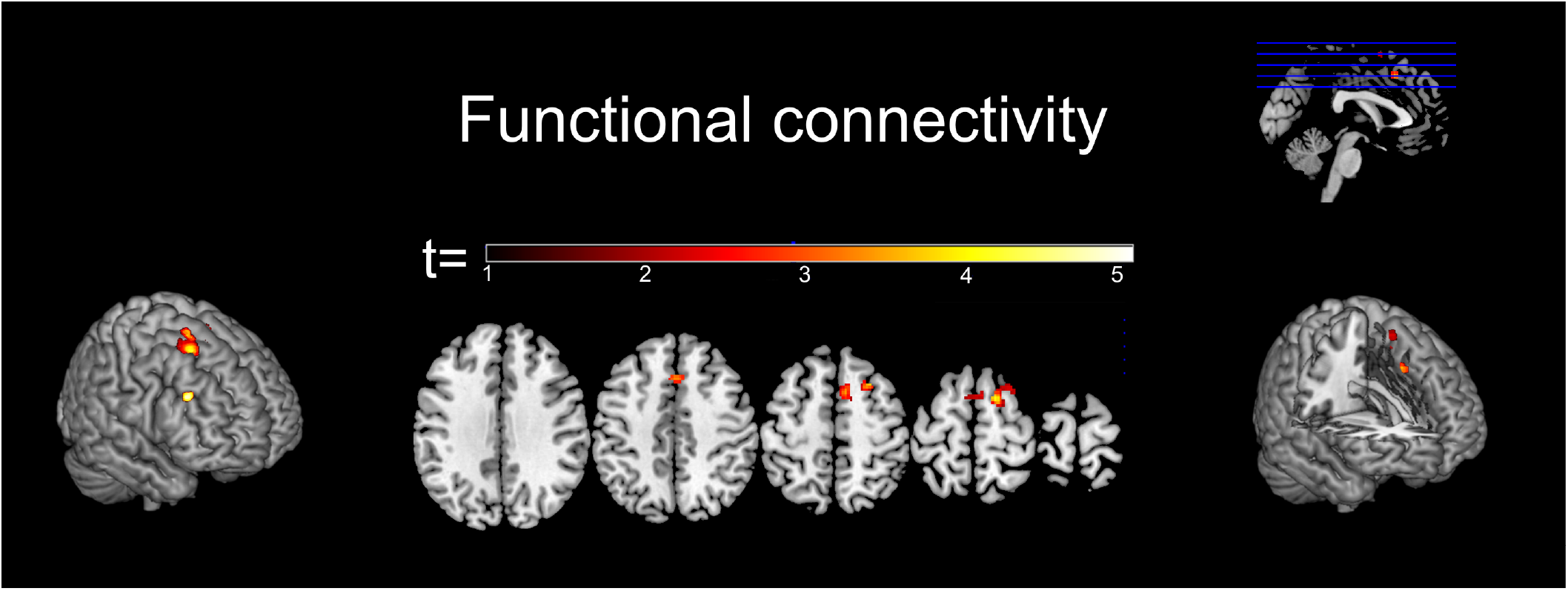
An increase in functional connectivity between the right dorsolateral prefrontal cortex and frontal/sensorimotor areas was observed after haloperidol/L-dopa treatment during N-back task performance (1-, 2-, and 3-back vs 0-back) (n=12).

**Table 4.** Brain areas in which haloperidol/L-dopa produced greater functional connectivity with DLPFC compared to placebo. Brodmann area (BA) and peak t-values, peak p-values and MNI coordinates indicate peak voxels within the clusters.

| General Area | Specific Area | BA | p-value<br>(FWE) | t-value | coordinates (MNI) |  |  |
| --- | --- | --- | --- | --- | --- | --- | --- |
|  |  |  |  |  | X | Y | Z |
| Frontal Cortex | R. Superior frontal gyrus | BA8 | 0.0174 | 4.41 | 24 | 16 | 58 |
|  | Bilat. Paracingulate gyrus | BA32 | 0.0353 | 4.06 | 0 | 20 | 46 |
|  | R. Middle frontal gyrus | BA45 | 0.0357 | 4.82 | 46 | 30 | 32 |
| Sensorimotor areas | R. Premotor cortex | BA6 | 0.0158 | 4.49 | 16 | 6 | 66 |
|  |  |  | 0.02 | 4.16 | 8 | 12 | 58 |
|  | L. Premotor Cortex | BA6 | 0.0387 | 3.14 | -2 | 6 | 66 |

## DISCUSSION

Here, we provide preliminary data for the effects of combined haloperidol/L-dopa administration on working memory-related brain activation. It was hypothesised that the combined haloperidol/L-dopa administration would induce changes in brain activity associated with N-back task performance, potentially by increasing SNR. In line with this hypothesis, smaller drug-induced differences in activity for the 2-back vs. 0-back contrast were observed in a large number of brain areas, most notably in the occipital/temporal cortex. Further, we observed drug-induced accentuated connectivity between DLPFC and right frontal brain areas (superior and middle frontal gyrus, and paracingulate gyrus) as well as sensorimotor areas (bilateral premotor cortices).

Specifically, the current results show increased functional connectivity of the DLPFC for *all* N-back versus 0-back conditions after haloperidol/L-dopa treatment_compared with placebo. An accumulating body of evidence suggests that dopamine agonism increases the strength of PFC-mediated networks that essentially underlie higher-order functions. For example, enhanced network integration during N-back performance was also observed after administration of the noradrenaline transporter inhibitor atomoxetine, which increases prefrontal catecholamine levels (Shine *et al*., 2017). Atomoxetine also enhances DLPFC activity during a response inhibition paradigm (Chamberlain *et al*., 2009) and enhances DLPFC functional connectivity during the N-back task (Hernaus *et al*., 2017). Importantly, L-dopa enhanced memory load-dependent working memory performance that coincided with increased frontal low theta power (Eckart *et al*., 2014).

D1 antagonism, however, has been reported *decrease* functional connectivity between DLPFC and premotor motor areas that were also identified in the current study (Rieckmann *et al*., 2012). Additionally, D2 blockade has been shown to decrease cortico-striatal connectivity in humans (Cole *et al*., 2013) and rats (Gass *et al*., 2013) at rest and decreases frontal cortical activity related to response inhibition (Luijten *et al*., 2013) and working memory (Goozee *et al*., 2016). Thus, similarities between our observation of drug-induced increase in connectivity within N-back network and other studies of (non-selective) dopamine agonism favour the interpretation that a combination of haloperidol and L-dopa may lead to D1-dependent accentuation of prefrontal cortex-mediated network integrity.

The drug-induced effects on occipital/temporal cortex activation were unexpected as these areas are not part of the typical task networks. Working memory-related activity in fronto-parietal regions during the placebo session aligned well with expectations from a meta-analysis of N-back studies (Owen *et al*., 2005), and did not include occipital/temporal cortex. Currently observed activity decreases in Brodmann areas 18 and 19 are part of the extrastriate cortex and are generally involved in processing of lower level visual information, such as motion, colour and contrast (Grill-Spector *et al*., 2004), while working memory-related activity changes in these regions are not typically observed during the N-back task. The results are also unlikely to be explained by simple changes in visual processing time as no reaction times differences were observed. However, drug-induced changes outside of the typical task network have been observed previously. Furey *et al*. (2000) showed that cholinergic enhancement can increase extrastriate cortex activation during encoding, which was accompanied by better working memory performance and reduced requirements for prefrontal activity. Therefore, even though the extrastriate cortex is not typically involved in specific working memory processes, drug induced modulation of its activation can coincide with changes in performance.

The presently observed decrease in occipital/temporal cortex activation may be a direct consequence of local D1 or D2 receptor modulation or indirect action via other regions (Yoon *et al*., 2006). Considering direct action, the presence of dopamine D1 and D2 receptors in this region is low compared with other cortical areas (Abi-Dargham *et al*., 2000) (www.brain-map.org), potentially limiting direct local effects. Importantly, functional imaging shows that dopaminergic modulation with L-dopa does not alter rCBF measured with positron emission tomography in the posterior occipital/inferior temporal areas (Hershey *et al*., 2003) and haloperidol reduced relative, but not absolute, blood flow in the posterior inferior temporal lobe in healthy volunteers given a single dose of 3mg (Handley *et al*., 2013). However, activity changes only occurred during the 2-back condition, and are thus unlikely to be explained by changes in blood flow, which would be expected to impact all conditions. In contrast, current evidence favours an indirect D2-receptor mediated decrease in task-related activation. Brassen *et al*. (2003) observed a decrease in visual stimulation-evoked BOLD response in similar visual areas (BA18/19) after intravenously-administered haloperidol. Gibbs *et al*. (2005) and Vytlacil *et al*. (2009) observed modulation of the visual association cortex activity in a working memory task after administration of the D2-agonist bromocriptine. Taken together, there is evidence that D2 receptors modulate working memory related occipital cortex activation, and those effects are likely to be the result of altered activation of regions connected to the occipital cortex.

As no significant effects on performance were observed, the behavioural consequences of combined haloperidol and L-dopa administration remain unclear. Despite the small sample size, the non-significance of the drug induced effects is in line with previous results suggesting that haloperidol 2 mg only has minor effects on cognitive performance, while higher doses from 3 mg do induce impairments (Legangneux *et al*., 2000). Alternatively, higher L-dopa doses may have resulted in enhanced N-back performance, perhaps represented by increasing response speed, without decreasing accuracy (Eckart *et al*., 2014).

A clear limitation of the present study is that it was designed as a proof-of-principle study to explore the potential effects of combined haloperidol/L-dopa on working memory-related brain activation. Therefore the study did not include all treatment conditions for a full factorial design and, thus, the contribution of L-dopa and haloperidol alone, as well as the interaction between treatments, could not be determined. Nonetheless, within the context of understanding combined treatment effects, we were able to confirm the utility of fMRI as a highly sensitive measure to such changes.

A final limitation concerns the number of included participants in this study. As data from only 12 participants were analysed any strong conclusions cannot be drawn from this data. For example, post-hoc calculations to determine achieved power showed that the main drug effect on reaction time could only be determined with power of 0.40. Nonetheless, we observed significant drug effects on brain activation that are in accordance with previous studies using dopamine manipulations, confirming effective manipulations.

The precise implications for disorders with working memory impairment such as schizophrenia are complicated by the fact that [18F]-DOPA imaging studies have consistently demonstrated increased dopamine synthesis capacity [e.g. (Howes *et al*., 2007)]. This would argue against the use of L-dopa, which would further elevate presynaptic dopamine function in the striatum. Studies in experimental animals combined with our findings point towards extra-striatal action of L-dopa combined with haloperidol as an important mechanism to improve cognitive performance, thus predicting utility for drugs that favour modulation of cortical dopamine projections. Therefore, in conclusion, the present study provides initial support for the combination of an indirect dopamine agonist with a dopamine antagonist having the required effects, but any strong conclusion should await replication in a larger study, including a full factorial design and higher doses of L-dopa

## Acknowledgements

The authors would like to acknowledge Prof. Dr. Shitij Kapur for insightful discussions. This research received no specific grant from any funding agency in the public, commercial, or not-for-profit sectors.

